# Curcumin-silk tyrosine crosslinked hydrogels

**DOI:** 10.1101/2025.02.14.638374

**Authors:** Aswin Sundarakrishnan

## Abstract

Systemic chemotherapy is still the first-line treatment for cancer, and it’s associated with toxic side effects, chemoresistance and ultimately cancer recurrence. Rapid gelling hydrogels can overcome this limitation by providing localized delivery of anti-cancer agents to solid tumors. Silk hydrogels are extremely biocompatible and suitable for anticancer drug delivery, but faster gelling formulations are needed. In this study, we introduce a rapid gelling hydrogel formulation (< 3 minutes gelling time) due to chemical crosslinking between silk fibroin and curcumin, initiated by the addition of minute quantities of HRP and H_2_O_2_. The novel observation in this study is that curcumin, while being a free-radical scavenger, also participates in accelerating silk di-tyrosine crosslinking in the presence of HRP and H_2_O_2_. Using UV-Vis, rheology and time-lapse videos, we convincingly show that curcumin accelerates silk di-tyrosine crosslinking reaction in a concentration-dependent manner, and curcumin remains entrapped in the hydrogel post-crosslinking. FTIR results show an increase in secondary beta-sheet structures within hydrogels, with increasing concentrations of curcumin. Furthermore, we show that curcu-min-silk di-tyrosine hydrogels are toxic to U2OS osteosarcoma cells, and most cancer cells are dead within short time scales of 4 hours post-encapsulation.

**Graphical abstract:** 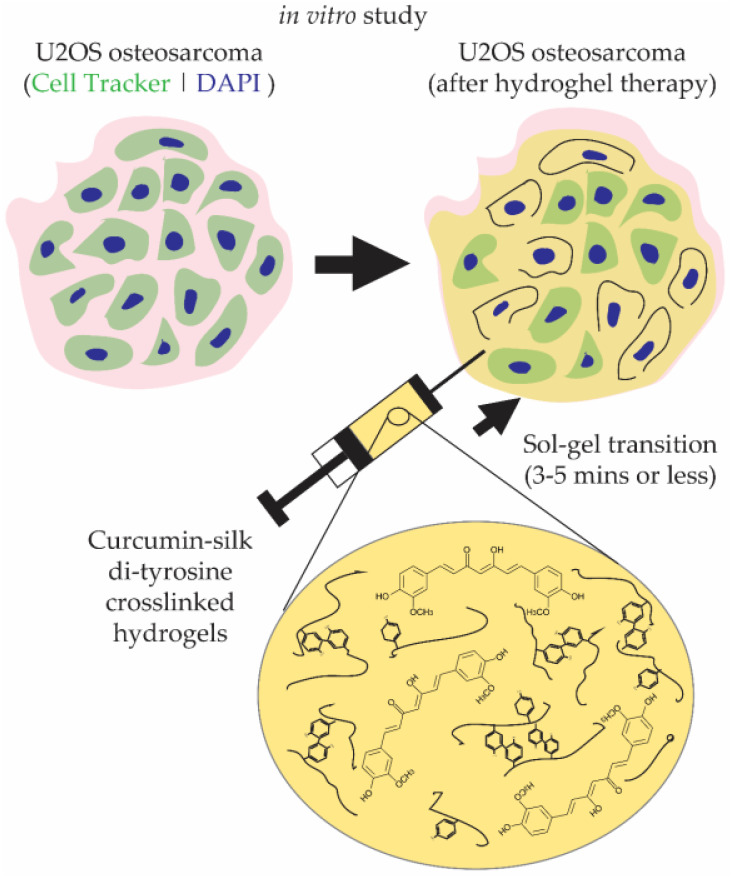

## 1. Introduction

Cancer is a significant healthcare burden, and it is the second leading cause of mortality in the United States, with ∼2 million new cases and 0.6 million projected deaths in 2025 [1]. Chemotherapeutics are still the first line of treatment for most solid tumors, and they are delivered intravenously, resulting in toxic side effects, causing chemoresistance, ineffective cancer killing, and ultimately leading to cancer recurrence. This is especially true for osteosarcoma patients, where recurrence rates are up to 40-50% even after aggressive systemic chemotherapy and surgery [2,3]. Therefore, targeted delivery of anticancer drugs is a much more effective strategy for killing solid tumors, especially prior to metastasis.

Hydrogel drug delivery vehicles are used for treating solid tumors, owing to their biocompatibility, mechanical properties, degradability and their ability to provide localized delivery of anti-cancer drugs over long periods of time [4,5]. Nanoparticles have also been widely used for anti-cancer drug delivery, but they are often sought out in conjunction with hydrogels as combinatory systems [6,7]. Smart thermoplastic hydrogels are often preferred because they undergo a quick sol-gel transition at 37°C and can be delivered in liquid form, filling the area surrounding tumors easily [8,9]. Silk fibroin protein hydrogels are well-suited for treating solid tumors, as they are smart-thermoplastics capable of gelling in vivo, with excellent biocompatibility. Silk fibroin hydrogels are better suited than other natural extracellular matrix hydrogels (e.g., collagen, gelatin, hyaluronic acid etc.), as their amino acid composition promotes degradability, but prevents cancer cell growth due to the lack of cell binding sites. There have been many previous studies utilizing silk fibroin hydrogel systems for anti-cancer drug delivery, however most of these studies do not support a quick gelling hydrogel formulation that can be delivered to the target tumor site [10-16]. Those studies that developed quick gelling hydrogels (<5 minutes gelling time), suffered from other limitations such as (1) high silk concentrations, which can be difficult to administer, or (2) were combinatory products, which can be difficult to translate in the clinic.

A review of recent literature shows that chemical crosslinking methods consistently delivered the quickest gelling silk hydrogel formulations. Yan et al. used minute quantities of horseradish peroxidase (HRP) and hydrogen peroxide (H_2_O_2_) to induce silk hydrogel gelation [11]. Rapid gelling hydrogels from 4-40 minutes were fabricated, but very high concentrations (∼16% w/v) of silk fibroin, and higher concentrations of HRP were required to achieve shorter crosslinking times. The fabricated hydrogels in this study were able to kill chondrogenic cancer cells in vitro and in vivo within 7 days, coinciding with beta-sheet formation. Li et al., fabricated silk hydrogels by the physical drying process, with or without curcumin, a well-known anti-cancer drug [12]. Interestingly, silk hydrogel gelation-time decreased with increasing concentration of curcumin, and hydrogel gelation times between 30 minutes to 5 hours are reported. Because the study was not focused on treating cancer, the authors did not investigate faster gelation times. In another study that utilized physical crosslinking methods, Chaala et al. prepared silk fibroin/hyaluronic acid composite hydrogels via sonication, and a sonication time of at least 5 seconds was required to initiate gelling, taking anywhere from 30 minutes to 2 hours to achieve complete gelation [13,14]. Peng et al. fabricated silk/PEG composite hydrogels with gelling times closer to 30 minutes [15]. These hydrogels could be mixed with polyvinylpyrrolidone iodine (PVP-I) for killing osteosarcoma cells both in vitro and in vivo. Another simple gelling formulation was developed by Laomeephol et al., by mixing silk fibroin with Dimyristoyl glycerophosphorylglycerol (DMPG)-based liposomes [16]. Hydrogels produced using this technique had extremely rapid gelation times of 3 minutes, but gelation effect was only observed with certain liposomal formulations. Hydrogels prepared with curcumin encapsulating liposomes displayed anti-cancer properties and killed MDA-MB-231 breast cancer cells.

Herein, we introduce a rapid gelling hydrogel formulation (<3-minute setting time) due to chemical crosslinking between silk fibroin and curcumin. Curcumin is a yellow-orange polyphenolic compound taken from the *Curcuma longa* plant that is known to have potent anti-cancer and anti-oxidant properties [17]. But curcumin has limited therapeutic efficacy because of its very low bioavailability, and poor ADME (Adsorption, Distribution, Metabolism, Excretion) properties [18]. A human Phase I clinical trial showed curcumin is non-toxic up to 8mg/day, but this dosage will only produce plasma concentrations of 0.5-1 µM within 1–2-hour post administration [19,20]. Through the new rapid gelling hydrogel formulation observed in this study, curcumin could have improved bioavailability, and it could be delivered over the long-term in the human body.

The novel observation in this study is that curcumin, while being a free-radical scavenger, also participates in accelerating silk di-tyrosine crosslinking in the presence of HRP and H_2_O_2_. Using UV-Vis and rheological characterizations, we show that curcumin participates in the silk di-tyrosine crosslinking reaction, and it also accelerates the rate of the chemical reaction at the tested concentrations (i.e., 50µM - 2mM). Hydrogels produced using this technique could have extremely rapid gelling times of 3 minutes or less, dependent on curcumin and substrate concentrations. Furthermore, we show that curcumin is immobilized within the hydrogel matrix, and does not leak post crosslinking, suggesting chemical bonding with the hydrogel network. Using FTIR, we show that hydrogels formed by this chemistry display increased beta-sheet structures with increasing concentrations of curcumin. Finally, we show that curcumin-silk crosslinked hydrogels are potent anti-cancer agents, and they are extremely toxic to osteosarcoma cells in vitro even at the lowest tested concentrations.

## 2. Results and Discussion

### 2.1 Curcumin participates in di-tyrosine crosslinking of silk fibroin hydrogels

To investigate whether UV-Vis spectroscopy could be used to study silk di-tyrosine crosslinking, wavelength sweeps were conducted on aqueous silk+HRP (S+H) and silk+HRP+curcumin (S+H+C). Neither of these solutions contained hydrogen peroxide (H_2_O_2_) which initiates the chemical crosslinking reaction. **Figure 1(a)** shows the absorption spectrum of silk+HRP (S+H) with two maxima, one below 300 nm and the other at 410 nm. This is consistent with previous reports showing absorption maxima for pure silk fibroin and HRP at 280 nm and 400 nm, respectively. [21]. Previous studies also show that curcumin has a strong absorption maxima at ∼430 nm [22]. Corroborating these results, we found curcumin to have a similar absorption maxima at ∼430 nm when co-mixed with 3% (w/v) aqueous silk fibroin and horsheradish peroxidase (HRP) (S+H+C in **Figure 1(a)**). The absorption maximum of pure di-tyrosine is reported at 315 nm [23,24]. Because silk fibroin, curcumin and HRP do not have overlapping absorption maxima at 315 nm, we decided to use UV-Vis time sweeps to monitor the progression of di-tyrosine crosslinking in silk fibroin hydrogels.

**Figure 1.**
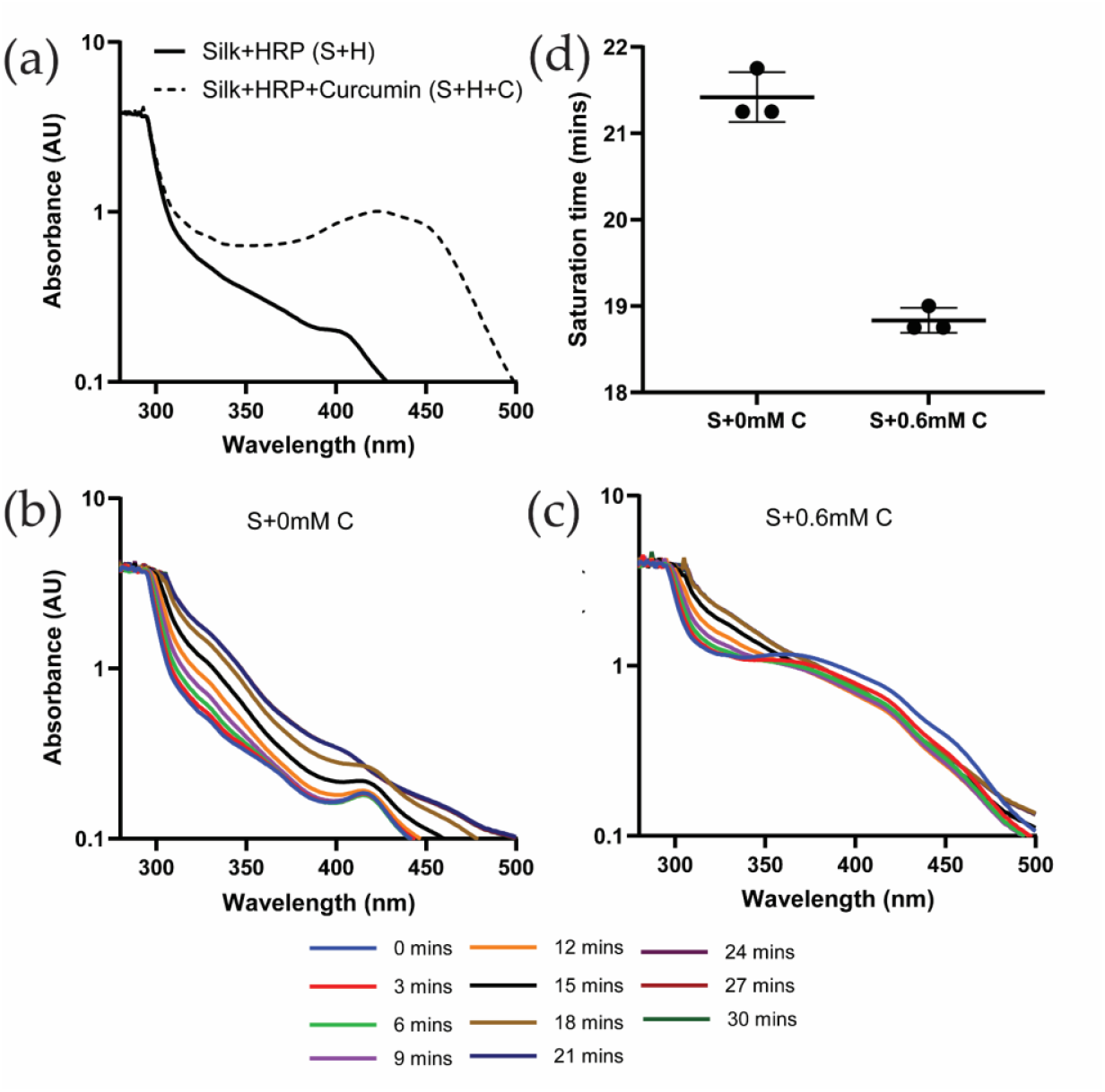
UV-Vis absorbance spectra of silk and curcumin-silk di-tyrosine crosslinked hydrogels: (a) UV-Vis absorbance of silk+HRP (S+H) and silk+HRP+curcumin (S+H+C) solutions without hydrogen peroxide (H_2_O_2)_ (b) UV-Vis wavelength and time sweeps of di-tyrosine crosslinked silk hydrogels; (c) UV-Vis wavelength and time sweeps of di-tyrosine crosslinked curcumin-silk hydrogels; (d) Saturation times of di-tyrosine crosslinked silk (S+0mM C) and curcumin-silk hydrogels (S+0.6mM C).

**Figures 1(b)** shows UV-Vis wavelength and time sweeps of silk di-tyrosine hydrogel gelation without curcumin over a time period of 30 minutes. To initiate gelation, minute quantities of hydrogen peroxide (H_2_O_2_) was added to the reaction mixture before recording the UV-Vis spectra in these experiments. The results are presented in a log-axis scale to better visualize the absorption bands across all wavelengths. In this experiment, UV-Vis spectra clearly captured the di-tyroine crosslinking reaction of silk fibroin as seen in **Figure 1(b)**. Variations in absorbance spectra were not observed when no H_2_O_2_ was added to the reaction mixture (results not shown). Saturation of the reaction was observed at 21 minutes, as subsequent absorption bands overlapped with each other and no further increase in absorbance was observed (**Figure 1(b)**). Interestingly, the absorption maximum at 415 nm reduced in intensity as the di-tyrosine crosslinking reaction neared completion, suggesting inactivation of HRP during the chemical reaction [25].

**Figure 1(c)** shows UV-Vis wavelength and time sweeps of silk di-tyrosine hydrogel gelation with 0.6 mM curcumin over a period of 30 minutes. Once again UV-Vis spectra clearly captrued the di-tyrosine crosslinking reaction. The addition of hydrogen peroxide (H_2_O_2_) shifted the UV-Vis minima and maxima to 338 nm and 360 nm, from 350 and 430 nm (i.e., absorbance maxima for the S+H+C reaction mixture in **Figrue 1(a)**). As the reaction progressed, the 315 nm di-tyrosine absorption value increased, while the absorption maxima at 360 nm decreased as a function of time. Saturation of di-tyrosine crosslinking reaction was observed at 18 minutes, as subsequent absorption bands overlapped with each other and no further increase in absorbance was observed (**Figure 1(c)**).

To better understand the effect of curcumin on di-tyrosine crosslinking dynamics, and to get more accurate estimates of di-tyrosine reaction saturation times, UV-Vis time sweeps were conducted at the 315 nm di-tyrosine absorbance. Results from these experiments are shown in **Figure 1(d)**, where di-tyrosine saturation times are compared between the two groups. As seen in the figure, addition of 0.6 mM circumin significantly (p<0.0002) reduced silk di-tyrosine crosslinking saturation time by ∼2.5 minutes. Taken together, we conclude that curcumin participates in the silk di-tyrosine crosslinking reaction, and reaction saturation is faster in the presence of curcumin.

The observation that curcumin promotes di-tyrosine crosslinking in the presence of HRP and H_2_O_2_ is a novel observation because curcumin is well-known anti-oxidant. Previous research has showcased that curcumin can act as a free radical scavenger when introduced into an environment containing hydrogen peroxide [26-29]. The anti-oxidant capacity of curcumin has been attributed to the diketone group and the two phenolic rings, but curcumin can promote anti-oxidant activity by inducing enzymes and related cell signaling pathways [26,27]. The anti-oxidant behavior of curcumin has primarily been observed at lower concentrations (<20 µm)[26,28], but at higher concentrations, this effect disappeared, and moderate cell-type specific cytotoxicity was observed [29]. It is possible that the rate promoting activity of curcumin observed in this study could be concentration dependent, although even at the lowest concentration of 50 µM, curcumin still promoted di-tyrosine crosslinking. Li et al., suggested that curcumin could bind to the hydrophobic domains of silk to accelerate silk hydrogel gelling, although we did not observe an increase in absorbance in S+H+C (silk+HRP+curcumin) reaction mixtures in the absence of H_2_O_2_ in shorter time scales (<30 minutes). The exact mechanism by which the curcumin promotes free-radical dependent di-tyrosine crosslinking requires further investigation, and this could have novel implications for its use as a therapuetic.

### 2.2 Curcumin accelerates silk di-tyrosine hydrogel gelation

To characterize the setting time, di-tyrosine crosslinking of hydrogels was characterized using rheology. **Figure 2(a)** shows rheological time sweeps of silk di-tyrosine crosslinking reaction with no curcumin added to the mixture (S+0 mM C). As per the results, the storage modulus (G’) starts to increase around 5 minutes and continues to increase until 20 mins, when the modulus reaches saturation. The loss modulus (G’’) was very low compared to the storage modulus (G’) throughout the crosslinking reaction, and no crossover was observed between the two. Crossover was previously utilized to study the setting time (gel-sol transformation) of polymer-based hydrogels [30]. However, there are limitations to this measurement, as crossing over was not observed in quick setting gels in some studies [31,32]. Furthermore, rheological time sweeps conducted on similar biomaterial hydrogels such as agarose, collagen and fibrin did not show crossing over behavior [31].

**Figure 2.**
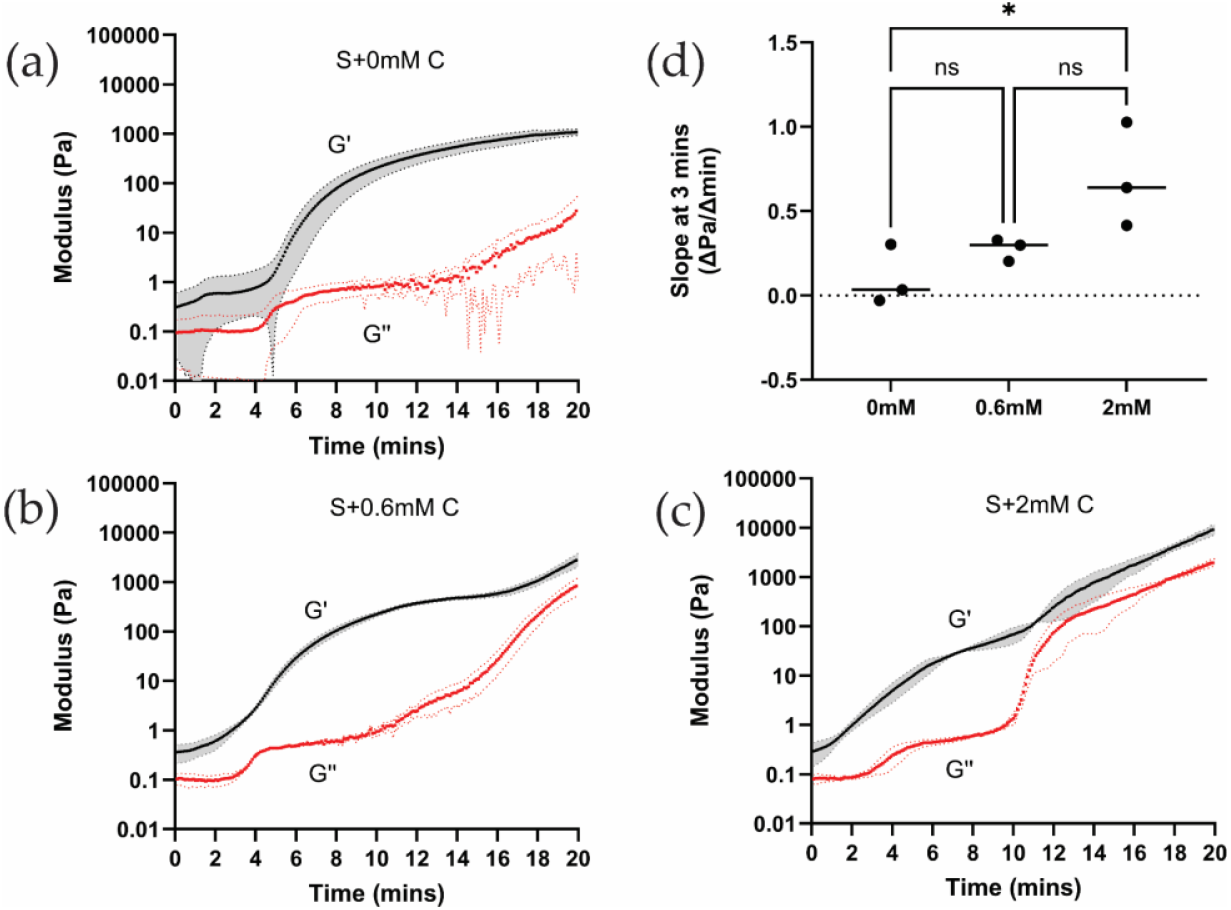
Rheology time sweeps of silk and silk-curcumin di-tyrosine crosslinked hydrogels: (a) Rheological time sweeps of silk di-tyrosine crosslinking hydrogels with no curcumin (b-c) Rheological time sweeps of di-tyrosine crosslinked silk hydrogels with 0.6mM and 2mM curcumin (d) Saturation times of di-tyrosine crosslinked silk and silk-curcumin hydrogels.

**Figure 2(b)** shows rheological time sweeps of silk di-tyrosine crosslinking containing curcumin (S+0.6 mM C). Addition of 0.6 mM curcumin caused the storage modulus (G’) to increase around 3 minutes, and it continues to increase up until 16 minutes where it reaches a pseudo-plateau point before entering a transition zone and increasing further. This is in contrast to silk di-tyrosine crosslinked hydrogels containing no curcumin (S+0 mM C), where the storage modulus starts to increase at 5 minutes and reaches saturation at 20 minutes, with no clear transition zone. Here again, the loss modulus (G’’) of the hydorgels remained well below the storage modulus (G’), and no crossing over was observed.

When the concentration of curcumin was increased to 2mM (S+2mM C), the storage modulus started to increase even earlier, between 0 to 2 minutes, reaching a pseudo-plateu point around 10 minutes, and then entering a transition zone and increasing further (**Figure 2(c)**). The rate of increase of the storage modulus is greater and occurs earlier in these hydrogels compared to the other hydrogel groups (i.e., S+ 0mM C and S+0.6 mM C). To quantitatively showcase this difference, the rate of change of storage modulus between 0-3 minutes was calculated for all three hydrogel groups. Comparing the intial reaction rates across all three hydrogel groups, a significant (p<0.0310) difference was observed between di-tyrosine crosslinking reactions of silk and silk-curcumin hydrogels. However, the reaction rates between the two groups containing increasing concentratons of curcumin were not significiant (**Figire 2(d)**). Based on these results we conclude that curcumin accelerates silk di-tyrosine crosslinking reaction rate upon initiation of the reaction.

To confirm that curcumin accelerates silk di-tyrosine crosslinking, a time-lapse video was captured with periodic inversion testing of hydrogel gelation ([Click here to view time-lapse video](https://1drv.ms/f/s!Avu8IfPbXNmmh75OOiSJ2ndG4csx3A?e=kf4WO2)). For this experiemnt, we tested 3 different concentrations of curcumin 0.05 mM, 0.6 mM and 2 mM. The hydrogels were crosslinked at 37°C, identical to the conditions used for the UV-Vis and rheological studies. The video was started immediately after adding H_2_O_2_ to all the groups, which is the start of the di-tyrosine crosslinking reaction. As seen in the video, the 2mM curcumin-silk di-tyrosine hydrogels (S+2mM C) set quickly, at approximately 2 minutes and 26 seconds. At this time, all the remaining hydrogels are still in liquid form, as seen in the video. With continued incubation at at 37°C, the setting time of other hydrogels groups could be approximated. Silk hydrogels containing 0.6 mM and 0.05 mM curcumin set at approximately 3 min 23 sec and 4 min 16 sec respectively. Silk hydrogels containing no curcumin had a setting time of approximately 5 minutes and 16 seconds. The video of the hydorgels undergoing phase change is appended to the online version of this article. The video clearly shows that hydrogels with curcumin have a much faster setting time compared to hydrogels without curcumin, once again confirming that silk di-tyrosine crosslinking reaction is accelerated by the addition of curcumin. A snapshot of the video showcasing the difference is included as **Figure 3(a). Figure 3(b)** confirms curcumin remained immobilized within the hydorgel matrix after di-tyrosine crosslinking, and no curcumin can be seen leaching out the hydrogel into water post-crosslinking.

**Figure 3.**
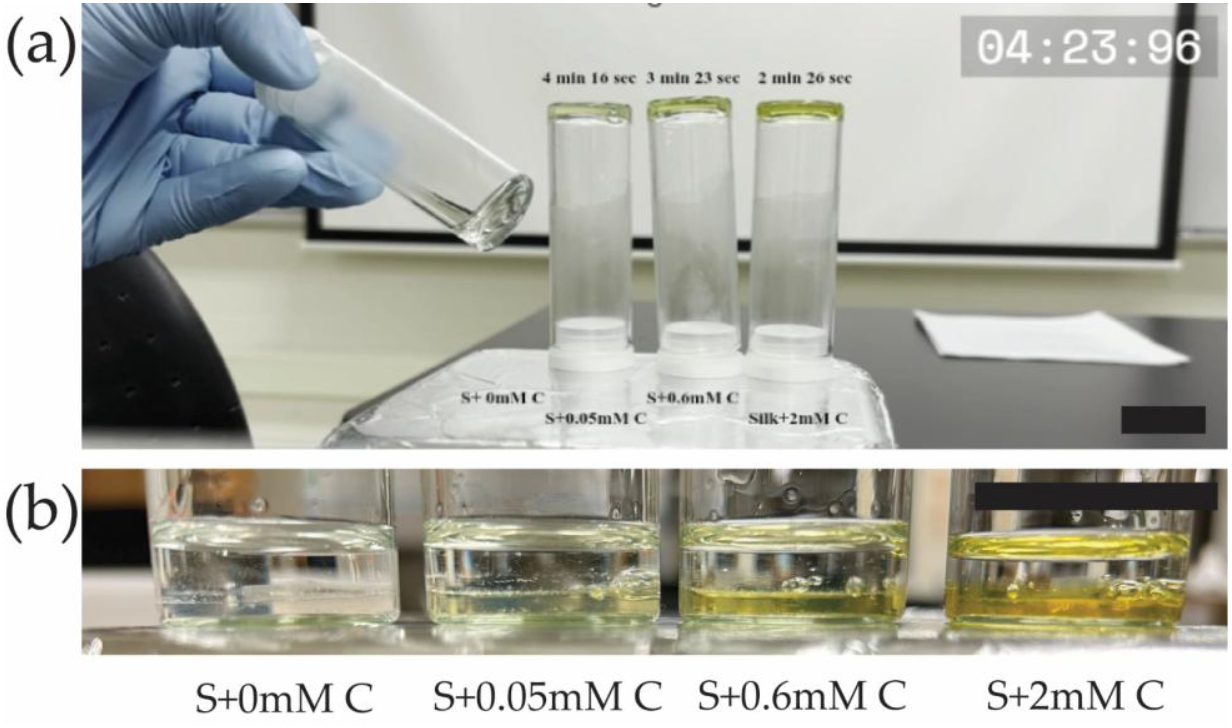
Curcumin-silk di-tyrosine crosslinking hydrogels (a) Snapshot of video [Click here to view time-lapse video](https://1drv.ms/f/s!Avu8IfPbXNmmh75OOiSJ2ndG4csx3A?e=kf4WO2) showing earlier setting times for silk di-tyrosine hydrogels containing curcumin compared to silk di-tyrosine hydrogels containing no curcumin, (b) Snapshot confirming curcumin is immobilized inside the hydrogels with no leakage into surrounding liquid. The scale bars are 1 cm.

### 2.3. ATR-FTIR spectra shows characteristic beta-sheet crystalline peaks in curcumin-silk dityrosine crosslinked hydrogels

Attenuated Total Reflectance Fourier-Transform Infrared Spectroscopy (ATR-FTIR) was used to study the confrmational states of silk fibroin following the sol-gel transition. For this experiment all samples were lyophilized at the same time and subjected to the same experimental conditions. Lyophilized silk fibroin (S) which did not undergo di-tyrosine crosslinking (no HRP, H_2_O_2_ or curcumin were added) was used as the reference, and compared to lyophilized silk di-tyrosine crosslinked hydrogels with or without curcumin (i.e., S+0mMC, S+0.05mMC, S+0.6mMC, and S+2mMC).

**Figure 4(a)** shows stacked ATR-FTIR spectra of the different treatment groups for comparison. Across all samples there are clear amide I, II and III absorption bands, each representing the characteristic vibration modes C=O stretching, N-H deformation, and C-N stretching/N-H bending respectively [33]. While the profiles are similar across samples, there are a few differences in the peaks observed between the samples. The amide I peak had higher wavenumbers for silk fibroin (S; 1635cm-1) and silk di-tyrosine hydrogels (S+0mM C; 1620cm-1), and amide I is used as a marker for studying the secondary structrure of silk fibroin proteins [33-35].

**Figure 4.**
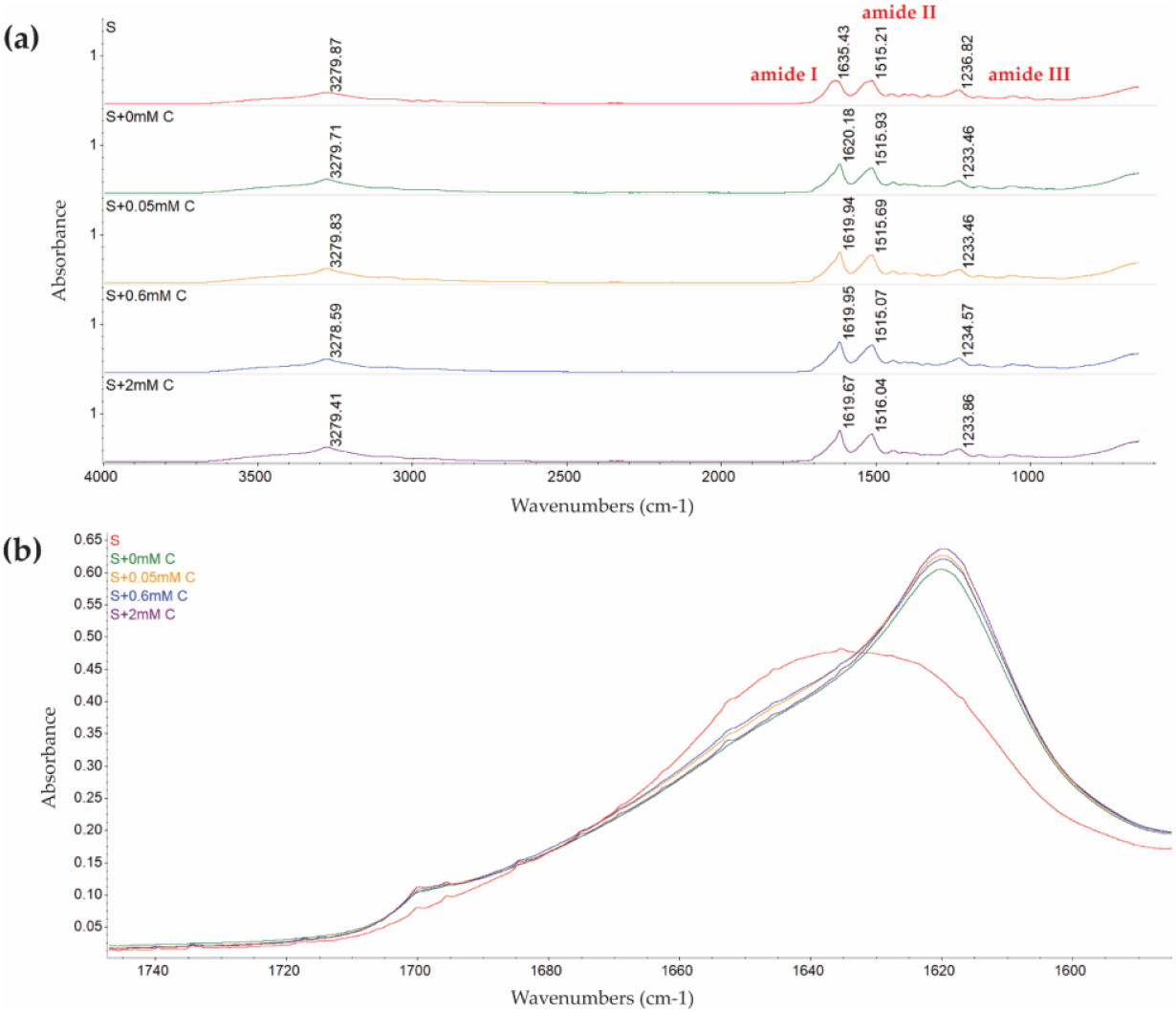
ATR-FTIR spectra of freeze-dried silk di-tyrosine hydrogels with or without curcumin, plotted as normalized absorbance versus wavenumber (a) Entire spectra between 650 cm-1 and 4000 cm-1, showcasing characteristic amide I, II and III silk fibroin peaks across all samples (b) Overlapped amide I band (1590-1720 cm-1) across all samples showcasing relative differences. All samples were freeze dried simultanously under identical experimental conditions. Legend: Silk fibroin (S); Silk di-tyrosine crosslinked hydrogels with no curcumin (S+0mM C); Silk di-tyrosine crosslinked hydrogel with 0.05mM curcumin (S+0.05mM C); Silk di-tyrosine crosslinked hydrogel with 0.6mM curcumin (S+0.6mM C); Silk di-tyrosine crosslinked hydrogel with 2mM curcumin (S+2mM C);

To further study the amide I band and understand relative differences across samples, the band is extracted from the spectra and presented separately in an overlapped fashion in **Figure 4(b)**. As seen from the figure, silk fibroin (S) has a relatively lower Amide I peak at 1635 cm-1, suggesting a mixture of beta-sheet and random coils, compared to dityrosine crosslinked silk hydrogels (i.e., S+0mMC, S+0.05mMC, S+0.6mMC, and S+2mMC), which display a strong peak at 1619 cm-1, corresponding with beta-sheet structures [34]. In addition to the characteristic 1619 cm-1 peak, silk di-tyrosine crosslinked hydrogels also displayed a strong shoulder peak between 1697-1703 cm-1, also indicating the formation of crystalline beta-sheet structures [34]. The peak height at 1619cm-1 and 1697cm-1, appeared to increase with increasing concentrations of curcumin, and curcumin-silk di-tyrosine hydrogels containing 2mM curcumin has the highest peak intensity in both these regions, while silk di-tyrosine hydrogels containing no curcumin (S+0mM C) had the lowest (**Figure 4(b)**). These results suggest that curcumin could increase the percentage of beta-sheet crystalline structures formed in silk di-tyrosine hydrogels post crosslinking. This would also explain the bi-phasic transition zone in rheology time sweeps, and higher storage moduli observed in curcumin-silk di-tyrosine hydrogels (**Figure 2(b-c)**). Further studies are required to dileneate the underlying mechanisms.

### 2.4. Curcumin-silk di-tyrosine crosslinked hydrogels are toxic to Ostoeosarcoma U2OS cancer cells

Curcumin is well known anti-cancer agent, and it has been investigated in clinical trials for many different types of cancers [36,37]. Curcumin has proven toxicity against osteosarcoma cells, but is limited by its poor bioavailability [38-40]. Because hydrogels can improve the bioavailabiltiy and also provide targeted delivery of curcumin, we studied the the effect of curcumin-silk di-tyrosine crosslinked hydrogels on encapsulated U2OS – osteosarcoma cells in an in vitro study. The cells were suspended in the hydrogel mixture pre-crosslinking and their viability was studied post-crosslinking using fluorescence concfocal microscopy.

Many previous studies have tested the effect of curcumin on cancers such as osteosarcomas, and toxicity was observed at dosages ranging from 10-100 µM [38,40]. It is important to note that curcumin was directly added to the media in most of these studies, unlike the current study where curcumin is encapsulated within the hydrogel matrix. In this study, we decided to test a larger range of curcumin dosages (0, 50 µM, 0.6mM and 2mM), as newer formulations of curcumin have been shown to have a bioavailability ranging from 30 nM – 2 mM, and dosages of 2g/day to 12g/day are well tolerated in human clinical studies [38,41].

**Figure 5 (a-d)** shows 3D fluorescence maximum projections of U2OS cells encapsulated within silk di-tyrosine hydrogels containing different concenrations of curcumin (0-2mM). Immunofluorescence signals from Cell Tracker-CMFDA dyes and DAPI clearly showcase a dose-response in viability across the different treatment groups within 4 hours of initial seeding. The 4-hour time point was chosen to allow sufficient time for cells to recover or undergo apoptosis. Very few viable cells (green) were observed within the 2mM curcumin treatment group (**Figure 5(d)**) at 4 hours. Silk di-tyrosine hydrogels containing no curcumin (S+0mM C) had the highest number of viable cells as seen in **Figure 5(a). Figure 5(e)** shows percentage of cellular viability across the different treatment groups, and treatment with curcumin, even at the lowest concentration of 50µM displayed significant toxicity (p<0.0008) compared to the control group. Almost all osteosarcoma cells were killed in the treatment group containing 2mM curcumin (<10% viable cells).

**Figure 5.**
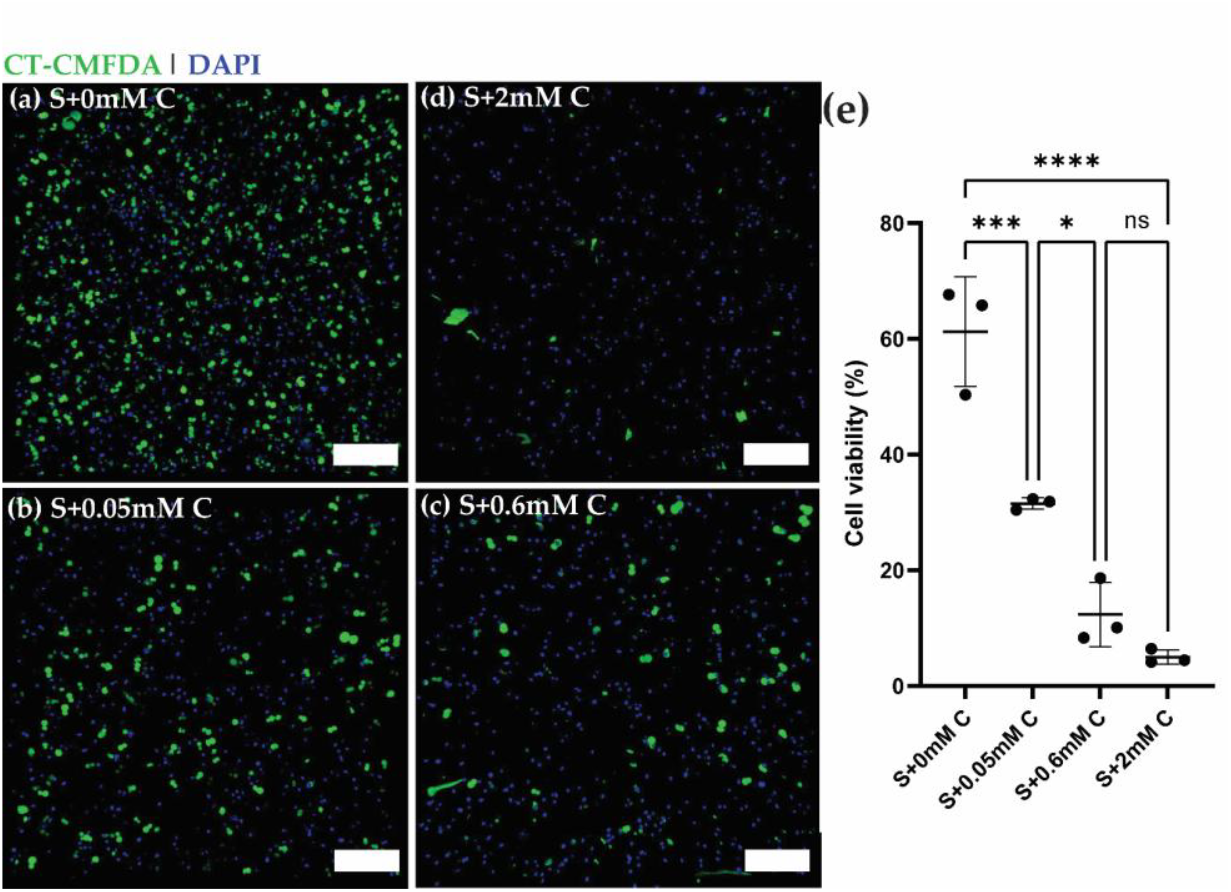
Viability of U2OS – Osteosarcoma cells encapsulated within curcumin-silk dityrosine hydrogels (a-d) Representative confocal 3D maximum projections of U2OS cells seeded within silk di-tyrosince hydorgels with different concentrations of curcumin (0, 0.05, 0.6 and 2 mM curcumin) 4 hours after cell seeding. All cells are stained with Cell Tracker-CMFDA green dye (live) and DAPI (dead). (e) Percent cell viability of osteosarcoma cells across different groups. Scale bars are 200 µm.

## 3. Conclusions

In summary, we convincingly show that curcumin accelerates silk di-tyrosine crosslinking reaction in the presence of HRP and H_2_O_2_. We show this rate promoting effect of curcumin using three different methodologies - UV-Vis spectroscopy, rheology and time-lapse videos **([Click here to view time-lapse video](https://1drv.ms/f/s!Avu8IfPbXNmmh75OOiSJ2ndG4csx3A?e=kf4WO2))**. To better understand the underlying mechanism, we performed FTIR, and this shows an increase in silk beta-sheet structures with increasing concentrations of curcumin. Finally, using fluorescence confocal microscopy we also show that curcumin-silk d-tyrosine crosslinked hydrogels are extremely toxic to U2Os-osteosarcoma cells in short time scales. While there are many previous studies on curcumin and silk, our study is the first in showcasing the rate-promoting activity of curcumin during di-tyrosine crosslinking. This is a novel observation because curcumin is also a well-known free radical scavenger, and one would expect curcumin to slow down di-tyrosine crosslinking. It is unclear why curcumin can accelerate a chemical reaction with hydrogen peroxide as the substrate. We postulate that this could be a concentration dependent effect, and it could be an important consideration while manufacturing curcumin-based therapeutics. Taken together, we showcase rapidly gelling curcumin-silk hydrogel formulations that can undergo sol-gel transitions in 3 minutes or less for anti-cancer drug delivery to solid tumors.

## 4. Materials and Methods

### 4.1. Preparation of aqueous silk solution

Aqueous silk solutions were prepared using well-established protocols reported earlier [42]. Briefly, 5g of *B. mori* silkworm cocoons were placed in 2L of boiling water containing 0.02 M Na_2_CO_3_ (Sigma-Aldrich, St. Louis, MO) solution for 30 minutes. The procedure is done to separate pure silk fibroin proteins from sericin. Subsequently, the silk fibroin fibers were washed three times in distilled water, and each wash step included a 15–20-minute soak time to remove the separated sericin protein. Air-dried fibers were then solubilized in 9.3M Lithium Bromide (LiBr; Sigma-Aldrich, St. Louis, MO) at 60°C for 4 hrs., followed by dialysis in 2L of distilled water (water changes at 1, 3, 6, 24, 36, and 48 hrs.) using a regenerated cellulose membrane (3.5kDa MWCO, Spectrum Laboratories, Rancho Dominguez, CA). The solubilized silk fibroin protein was then centrifuged at the maximum setting to remove any particulate waste material from the silk fibroin solution. Silk fibroin solutions were filtered using a 0.2 µm filter for sterilization prior to use in cell culture experiments.

### 4.2. Preparation of curcumin-silk and silk di-tyrosine hydrogels

Silk di-tyrosine hydrogels were prepared as per previous protocols [35,43,44]. For di-tyrosine crosslinking silk fibroin, 10 µl of 1,000U/ml stock solution of HRP (Horseradish peroxidase type IV; Sigma-Aldrich, St. Louis, MO) and 10 µl of 165mM hydrogen peroxide (H_2_O_2;_ Sigma Aldrich, St. Louis, MO) were added to 1 ml of 3% (w/v) silk solution (final concentration of 10U/ml of HRP and 1.65mM H_2_O_2_) at 37°C. Silk fibroin can change mechanical properties depending on extraction procedure, silk fibroin type, and other variables such as storage time post-extraction etc. Therefore, while 3% (w/v) is suggested as a guide to repeat these experiments, silk concentration should be calibrated each time, to identify a working concentration for your target application and experiments. Curcumin (Sigma Aldrich, St. Louis, MO), the yellow-colored polyphenol found in *Curcuma longa* was utilized to prepare curcumin-silk di-tyrosine crosslinked hydrogels. Curcumin solubilized in pure ethanol was co-mixed at appropriate volumes with silk fibroin to achieve final curcumin concentrations of 0.05mM, 0.6mM, and 2mM, respectively. To initiate crosslinking, HRP and H_2_O_2_ were added at the concentrations described above, followed by incubation at 37°C.

### 4.3. UV-Vis Spectroscopy

UV-Vis measurements were made using an Agilent Technologies (Agilent Technologies, Santa Clara, USA) Cary 60 UV-Vis spectrophotometer fitted with a Quantum Northwest (Quantum Northwest, Liberty Lake, WA) TC125 Peltier unit for temperature control. The same protocol was followed for all UV-Vis experiments. Prior to each experiment, the Peltier unit was set to 37°C, and we waited until the cuvette holder reached the target temperature. A blank was performed using water at the beginning for baseline correction. One point five milliliters (1.5 ml) of silk di-tyrosine crosslinking mixture with or without curcumin was added to disposable polystyrene cuvettes and placed inside the instrument. Wavelength time sweeps recorded absorbance at wavelengths 250-500 nm at 1 nm increments. Wavelength and time sweeps repeated the wavelength sweep experiment every 3 minutes for 30 minutes, or until saturation was observed. Results are plotted as a function absorbance versus wavelength.

### 4.4. Rheology

Rheological measurements were performed on silk and curcumin-silk dityrosine crosslinked hydrogels using a TA Instruments Discovery HR-2 rheometer (Waters Corporation, Milford, MA). The setup included a 40 mm stainless steel conical plate (angle: 0.0349 rad) and a temperature-controlled Peltier plate set to 37°C. Briefly, silk or curcumin-silk di-tyrosine crosslinking mixture was prepared by mixing 600 µl of the different constituents, at the concentrations mentioned above, and 420 µl of the mixture was dispensed on the tester. Immediately, the cone geometry was lowered to the trimming gap for trimming excess liquid and then lowered to the specified final testing gap. Low viscosity silicone oil was placed around the outside edge of the cone geometry to prevent water evaporation. Dynamic time sweeps were conducted as per previously established protocols [35,43], at a frequency of 1Hz (6.283 rad/sec) and 1% strain until the sample reached a plateau modulus. Preliminary experiments showed that storage moduli of silk di-tyrosine crosslinked hydrogels reached a plateau within 20 minutes, and so all subsequent experiments were run for 20 minutes.

### 4.5. Time-lapse video

To study the setting time of silk and curcumin-silk hydrogels, time lapse videos were recorded. Hydrogels were placed inside glass vials on a hot plate set to 37°C. Videos were recorded once H_2_O_2_ was added to the hydrogel mixture. Setting time of hydrogels was studied by conducting a partial inversion test at regular time intervals. Upon visual observation, if the hydrogels remained in a solution state, they were placed back onto the hotplate to continue gelation. If the hydrogels appeared to have undergone sol-gel transition, they were inverted, and the time of inversion was recorded as their setting time. All samples were tested at each point to enable relative comparison of setting times.

### 4.6. Fourier Transform Infrared (FTIR) Spectroscopy

Silk fibroin (S) without HRP, H2O2 or curcumin, and silk di-tyrosine crosslinked hydrogels with or without curcumin (i.e., S+0mMC, S+0.05mMC, S+0.6mMC, and S+2mMC) were frozen at -80°C and then placed inside a custom-made freeze-drying apparatus. A vacuum pump and cold trap were set up to keep the pressure and temperature well below the triple point of water. The freeze-dried silk fibroin (S) served as the positive control, as the FTIR spectra for lyophilized silk fibroin is well-established by other studies [45]. Attenuated Total Reflectance Fourier-Transform Infrared Spectra (ATR-FTIR) were recorded using a Nicolet iS50R FTIR spectrometer (Thermo Fisher Scientific, Waltham, MA, USA), using previously well-established protocols [33,45]. The spectrometer was equipped with a single bounce diamond attenuated total reflectance (ATR) module with a refractive index of 2.4 and an active sample area diameter of 2 mm. Spectra of each of the samples were acquired by pressing the samples on the ATR crystal. Air was used as the reference and automatically subtracted from sample readings. Samples were analyzed in the frequency range from 650 – 4000 cm-1, with each measurement adding 25 interferograms at a resolution of 1 cm-1. The amide 1, amide 2 and amide 3 regions were identified based on wavenumber ranges specified in previous publications [45]. The presence of secondary structures, including random coils, alpha-helices and beta sheets were also based on wavelength numbers within the amide I region [34].

### 4.7. Cell culture and cellular viability staining

Human osteosarcoma cell line (U-2 OS; HTB-96; ATCC) was cultured in DMEM/F12 medium (Life Technologies, Grand Island, NY) containing 10% fetal bovine serum (FBS) and 1x penicillin/streptomycin (Penn/Strep). A master stock of U2OS cell line was obtained from ATCC, and experiments were conducted using early passaged cells. For viability experiments, U2OS cells were co-mixed with silk and curcumin-silk di-tyrosine hydrogel precursor solutions at a concentration of 1×106 cells/ml. Cell laden hydrogel precursor solutions were then incubated at 37°C until hydrogel gelation, and warm cell culture medium was quickly added to the cell laden hydrogels. Cellular viability and distribution were studied by incubating cell laden hydrogels in 10 µM of CellTracker Green CMFDA (CT-CMFDA) in serum free media for 20 mins at 37°C. Hydrogels were then fixed using 4% paraformaldehyde (PFA) in 1xPBS for 30 mins and washed 3X in 1xPBS. Subsequently, hydrogels were stained with nuclear stain DAPI (4’,6-diamidino-2-phenylindole, dihydrochloride; Life Technologies, Grand Island, NY) at a final concentration of 1 μg/ml. While all cells are stained with DAPI, only live cells retain the CT-CMFDA dye.

### 4.8. Confocal Microscopy

Cell Tracker CMFDA and DAPI stained cell laden hydrogels were imaged using a Nikon A1R MP confocal microscope equipped with a tunable laser. Excitation lasers and filters were chosen to enable detection of fluorescent emissions for CT-CMFDA (MW: 464.9 g/mol ex: 492 nm; em: 517 nm) and DAPI (MW: 320 g/mol; ex: 364 nm; em: 454 nm). Using the Z-stack feature, image stacks of 500 µm depth were acquired using a 10x objective. Step size was set to 30 µm for all groups based on the average cell size. Each sample was imaged and a total of 3 samples/group was used for achieving statistical significance. Images obtained from the green and blue channels were analyzed using ImageJ software. After an initial thresholding step, the Analyze Particles feature was used to count the total number of cells in each z-stack across all groups. The percentage of viable cells in each group was obtained from this data and plotted for each treatment group. Fluorescence 3D maximum projection images with both fluorescent channels were visualized using the 3D Viewer plugin in Fiji ImageJ for relative comparison.

### 4.9. Statistical analysis

Data are expressed as mean, and error bars denote standard deviation. The unpaired student’s t-test was used for comparisons of two groups. For experimental comparisons of more than two groups, two-way analysis of variance (ANOVA) and Tukey post-hoc analysis were used to determine statistically significant differences. Statistical significance was accepted at the p < 0.05 level and indicated in figures as * p <0.05, ** p <0.01, *** p <0.001, **** p <0.0001.

### 4.10. Editing

Microsoft Co-pilot integrated with Word and Office 365 was used to edit this article. Co-pilot was used to (1) generate a list of acronyms (2) identify and correct typos, and (3) identify and correct grammatical errors in the manuscript. The article writing and data interpretation were done by the author based on subject matter expertise.

## Supporting information

Time-lapse video of gelation

## Abbreviations

The following abbreviations are used in this manuscript:

ADME: Adsorption, Distribution, Metabolism, Excretion
ANOVA: Analysis of Variance
ATR-FTIR: Attenuated Total Reflectance - Fourier Transform Infrared Spectroscopy
CMFDA: 5-chloromethylfluorescein diacetate
DAPI: 4’,6-diamidino-2-phenylindole, dihydrochloride
DMEM/F12: Dulbecco’s Modified Eagle Medium/Nutrient Mixture F-12
FBS: Fetal Bovine Serum
FTIR: Fourier Transform Infrared Spectroscopy
HRP: Horseradish Peroxidase
H2O2: Hydrogen peroxide
LiBr: Lithium Bromide
MWCO: Molecular Weight Cut Off
Na2CO3: Sodium carbonate
PBS: Phosphate Buffered Saline
Penn/Strep: Penicillin/Streptomycin
PFA: Paraformaldehyde

## Author Contributions

This work was inspired by previously published work. A.S. conceptualized the project, and carried out the experiments, data analysis and writing for this project. A.S. did get help from undergraduate students who are acknowledged in the manuscript. A.S. has read and agreed to the published version of the manuscript.

## Funding

Experiments and research work was funded by the Start-up Faculty grant provided through the Materials Science & Biomedical Engineering Resource Center at the University of Wis-consin Eau Claire.

## Data Availability Statement

The raw data supporting the conclusions of this article will be made available by the authors on request.

## Acknowledgments

The author thanks undergraduate students Andres Jimenez, Trixie Tah and Seth Waalen for their technical help with experiments and lab work related to this project.

## Conflicts of Interest

The authors declare no conflicts of interest.

## Notes

https://1drv.ms/f/s!Avu8IfPbXNmmh75OOiSJ2ndG4csx3A?e=kf4WO2

### Competing Interest Statement

The authors have declared no competing interest.

